# Long-read isoform sequencing reveals tissue-specific isoform expression between active and hibernating brown bears (*Ursus arctos*)

**DOI:** 10.1101/2021.07.13.452179

**Authors:** Elizabeth Tseng, Jason G. Underwood, Brandon D. Evans Hutzenbiler, Shawn Trojahn, Brewster Kingham, Olga Shevchenko, Erin Bernberg, Michelle Vierra, Charles T. Robbins, Heiko T. Jansen, Joanna L. Kelley

## Abstract

Understanding hibernation in brown bears (*Ursus arctos*) can provide insight into many human diseases. During hibernation, brown bears experience states of insulin resistance, physical inactivity, extreme bradycardia, obesity, and the absence of urine production. These states closely mimic human diseases such as type 2 diabetes, muscle atrophy, renal and heart failure, cachexia, and obesity. The reversibility of these states from hibernation to active season allows for the identification of novel mediators with possible therapeutic value for humans. Recent studies have identified genes and pathways that are differentially expressed between active and hibernation seasons. However, little is known about the role of differential expression of gene isoforms on hibernation physiology. To identify both distinct and novel mRNA isoforms, we performed full-length RNA-sequencing (Iso-Seq) on three tissue types from three individuals sampled during both active and hibernation seasons. We combined the long-read data with the reference annotation for an improved transcriptome and mapped RNA-seq data from six individuals to the improved transcriptome to quantify differential isoform usage between tissues and seasons. We identified differentially expressed isoforms in all study tissues and showed that adipose has a high level of differential isoform usage with isoform switching, regardless of whether the genes were differentially expressed. Our analyses provide a comprehensive evaluation of isoform usage between active and hibernation states, revealing that differential isoform usage, even in the absence of differential gene expression, is an important mechanism for modulating genes during hibernation. These findings demonstrate the value of isoform expression studies and will serve as the basis for deeper exploration into hibernation biology.

## Background

Hibernation in bears has long been championed as a promising system for understanding the extremes of mammalian physiology and for identifying novel therapeutic targets [1, 2]. Annual hibernation in brown bears (*Ursus arctos*) involves massive physiological shifts to conserve energy during the food-scarce winter [3], and every organ system in bears demonstrates a suite of adaptations driven by the needs of hibernation. Hibernating bears exhibit certain phenotypes present in human disease, but importantly, these phenotypes do not themselves negatively impact the bears’ overall health [4-7]. For example, heart rate slows to 10 to 15 beats per minute [8], yet bears do not develop typical dysfunction that would be characteristic of severe bradycardia in humans. Bears prevent congestive heart failure or ventricular dilation by decreasing atrial contractility and increasing atrial and ventricular stiffness [8, 9]. Hibernating bears are also well known for maintaining muscle strength, morphology, and composition during hibernation in the near complete absence of weight-bearing activity [10, 11]. Bears also exhibit insulin resistance during hibernation but are insulin sensitive during the active season [7]. Although humans do not hibernate, the unique metabolic adaptations that evolved in hibernators could provide clues to develop new treatments for human metabolic diseases, such as obesity and type 2 diabetes [2]. In fact, the need to accumulate tremendous amounts of fat and the development of insulin resistance evolved in bears as a survival strategy [12-14]. Recent studies have shown that these hibernation-induced physiological shifts are associated with massive changes in the regulation and expression of thousands of genes across hibernation-relevant processes [15-17]. Notably, many of these genes are involved in complex metabolic and cellular signaling pathways (e.g., insulin signaling, metabolism) that play critical roles in a variety of biological processes across vertebrates, including humans [15, 18].

Over ten thousand genes have been shown to be differentially regulated in adipose, liver, and muscle tissues between active and hibernating states, providing a set of candidate genes involved in the regulation of cellular and physiological processes that underly the metabolic suppression observed in hibernation [15]. The genes identified represent key genes and genomic regions for testing hypotheses related to the evolution and regulation of hibernation. While it is known that mRNA isoforms vary between tissues, cell types, and developmental stages [19, 20] and play a role in cellular processes, studies in bears have focused on global gene expression levels have not investigated the mRNA processing shifts that result in different isoforms and thus proteome output that occur with active and hibernating seasons.

We hypothesize that different transcript isoforms contribute to the reversible states achieved during hibernation. Indeed, in brown bears and Himalayan black bears (*Ursus thibetanus ussuricus*), it has been shown that the amount of titin does not differ between active and hibernation seasons but the relative abundance of two prominent isoforms may explain increased ventricular stiffness during hibernation [21, 22]. However, isoform differences have been explored only in a few select cases and little is known about the role different isoforms contribute to the hibernation phenotype on a large scale. Because of the capability of sequencing full-length RNA transcripts, SMRT Sequencing is ideal for identifying the isoforms that are differentially expressed between seasons. We compare full-length isoforms between hibernating and active bears in three metabolically active tissues – skeletal muscle, liver, and adipose.

## Results

### High Correlation between Long- and Short-Read Data at Gene Level

We analyzed RNA-sequencing data from bears in active or hibernation seasons on three distinct tissue types (muscle, adipose, liver). Three of the bears were sequenced using the PacBio Iso-Seq protocol for full length RNA transcripts, and all six bears were sequenced using the short-read Illumina RNA-seq approach (Figure 1). The RNA-seq data was previously analyzed in [15]. We first combined all the long-read Iso-Seq data across samples and replicates and obtained a total of 6.1 million full-length HiFi reads (Table S1). After running the HiFi reads through analysis, mapping to the reference genome, and filtering for library artifacts, we obtained 76,071 unique, full-length isoforms ranging from 150 basepairs (bp) to 16.5 kilobases (kb) (mean: 3.2 kb). We then evaluated the gene-level correlation of long- versus short-read data. For this correlation, we used only transcripts that were present in both Iso-Seq and RNA-seq datasets. When comparing data from the same individuals (albeit sampled in different years), the highest correlations are within data type, regardless of season (active or hibernation) (Figure 2). There is also a high correlation between data types at the gene level, especially within the same tissues (Figure 2a). The lower concordance at the isoform-level within the Iso-Seq samples as compared to the within RNA-seq samples, is likely explained by the lower sequencing coverage of the long-read dataset (Figure 2b). Nevertheless, we see consistent gene-level correlation for the matching samples across data types, while samples from the same tissues or animals have higher correlation than different tissues, as expected.

**Figure 1.**
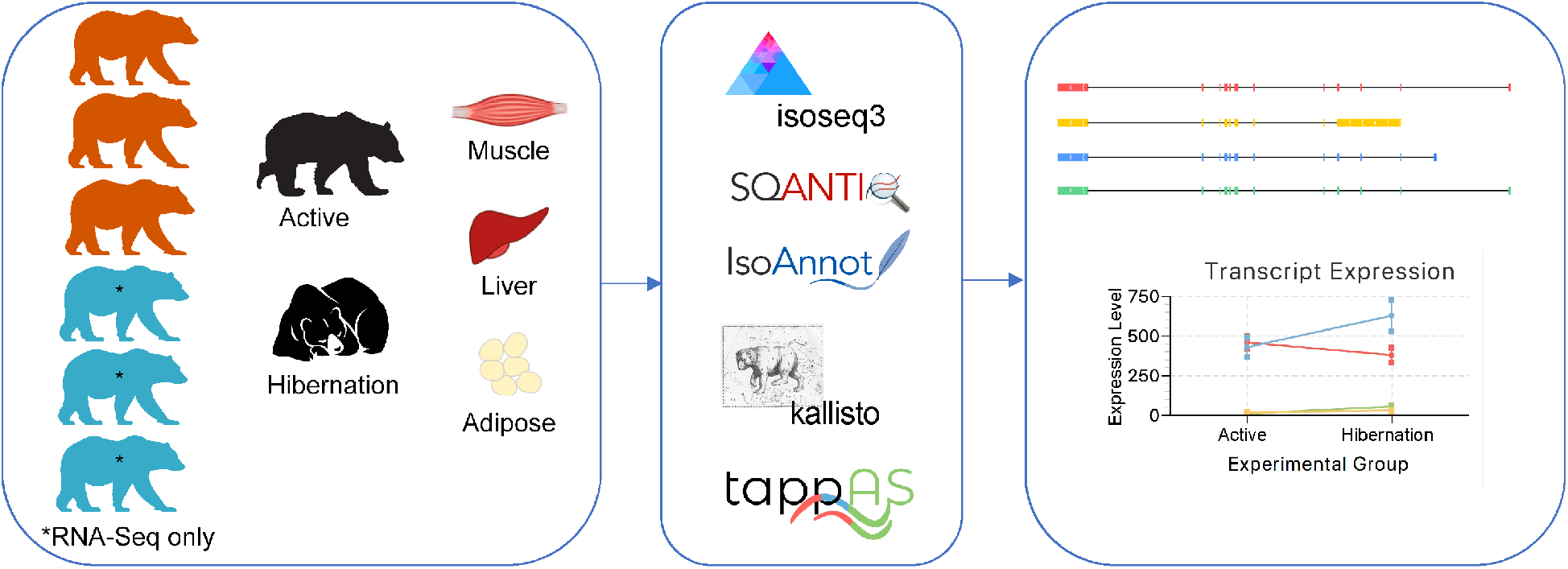
Bear transcriptome workflow. For each of the six bears, tissues (muscle, liver, adipose) were extracted during active or hibernation seasons. PacBio Iso-Seq and Illumina RNA-seq data were collected from three of the bears (orange) and Illumina RNA-seq data was collected from three additional bears (blue). Iso-Seq data was processed through a pipeline of isoseq3, SQANTI3, and IsoAnnot, before merging with the reference transcriptome to create a merged annotation set to map RNA-seq data using kallisto. Differential isoform expression and usage was determined using tappAS.

**Figure 2.**
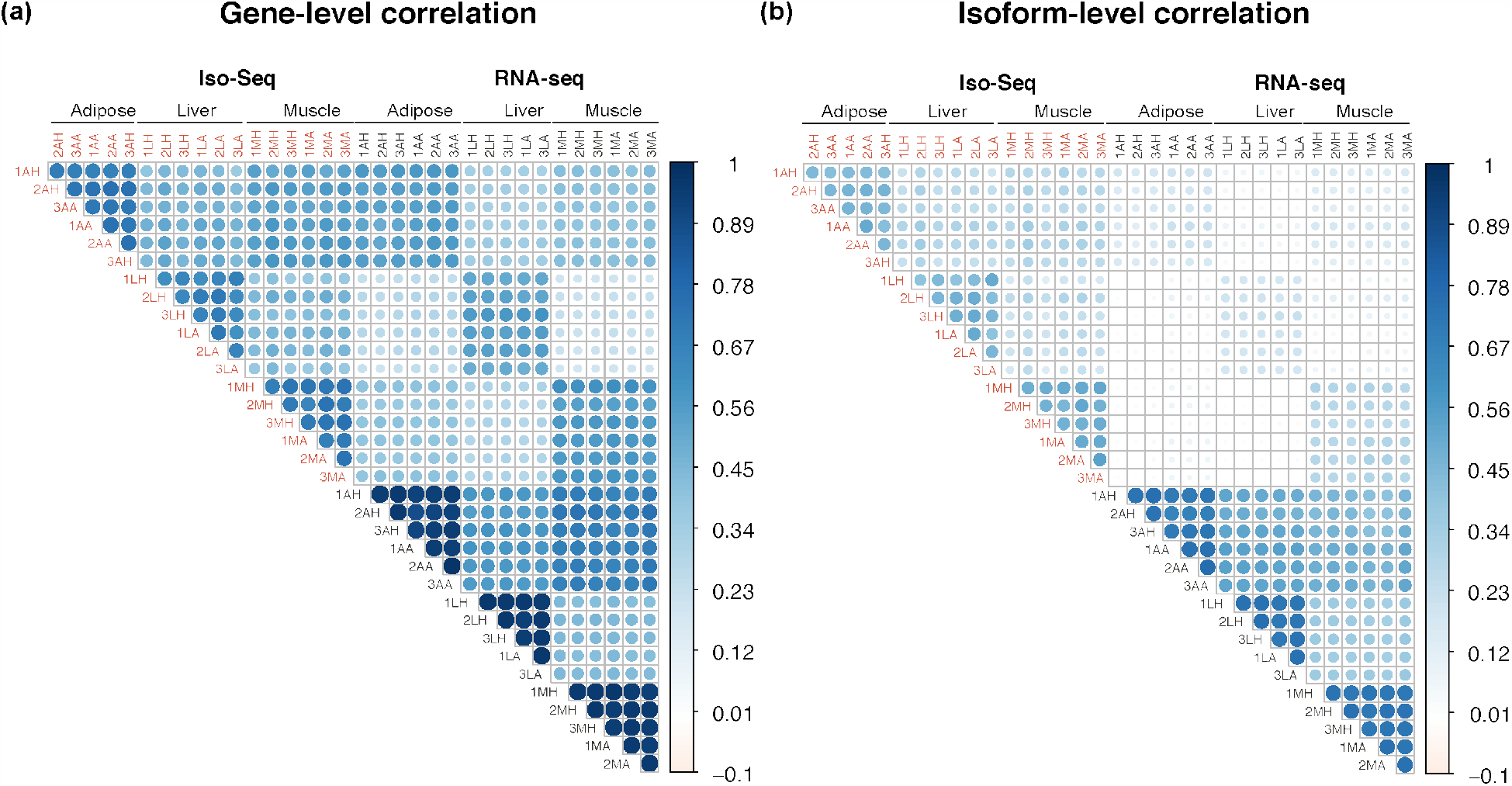
Gene- and isoform-level correlation between Iso-Seq and RNA-seq count data (log counts per million). Correlation measured with Pearson correlation. Samples are coded using three letters representing [animal], [tissue], and [season]. The three bears were numbered. The tissues were adipose (A), liver (L), and muscle (M). The seasons were hibernation (H) or active (A).

### An Improved Reference Transcriptome and Full-Length Isoform Annotation using Iso-Seq

The existing reference transcriptome contained 30,263 genes encompassing 58,335 transcripts. The Iso-Seq transcriptome was classified against the reference transcriptome and found to contain 12,018 known and 907 novel genes (Table S2). Compared with the reference annotation, 27.8% of the Iso-Seq isoforms were categorized as full-splice matches (FSMs; perfect matches to a reference transcript), while over half of the isoforms were novel isoforms (NIC, novel in catalog or NNC, novel not in catalog). More than 30% of the genes had complex splicing events (greater than six isoforms). Novel isoforms had a higher proportion of having a predicted nonsense mediated decay (NMD) effect (see the left panel of Figure S1a-S1d).

We merged the existing reference transcriptome with the new Iso-Seq transcriptome data, resulting in a total of 31,829 genes encompassing 107,649 transcripts. The merged data set had a reduced number of incomplete splice matches (ISM) and novel isoforms (NIC, NNC) while greatly increasing the number of known transcripts (Table S3), suggesting a more comprehensive representation of the transcriptome. When analyzing transcripts expressed in each tissue, by mapping RNA-seq data from six bears, we found that each tissue (muscle, liver, adipose) had different distributions of transcript structural changes, with adipose and liver having similar distributions and muscle having a much larger number of full-splice match (FSM) and fewer novel in catalog (NIC) transcripts (Figure S2). Importantly, the improved reference transcriptome, with the full-length transcripts originating from samples of interest, provides a starting point for the discovery of differential isoform usage (DIU) that would otherwise be missed. One example of such a finding is the isoform expression of the *CA2* gene (Figure S1e), described in more detail in later sections.

To evaluate the degree to which the Iso-Seq dataset captures the expressed transcripts in the tissues of interest, we mapped the ribosomal RNA-depleted (Ribo-Zero) short-read RNA-seq data of the same tissues (adipose, liver, muscle) and seasons (active and hibernation) to either a PacBio-only transcriptome, the reference-only transcriptome, a PacBio-reference merged transcriptome, or the reference genome (Figure 3). Given the lower sequencing depth of the PacBio data, we were not surprised to see a higher mappability of the short reads to the reference transcriptome than to the PacBio-only transcriptome. However, the merged PacBio-reference transcriptome showed the highest mappability of all transcriptomes. Of note, in the adipose and liver tissues, a higher proportion of intronic reads in the ribosomal-depleted RNA-seq data resulted in a much higher mappability using the reference genome. With the demonstrated improvement of the transcriptome by adding the Iso-Seq data, the merged (PacBio and reference) transcriptome was used for further analyses.

**Figure 3.**
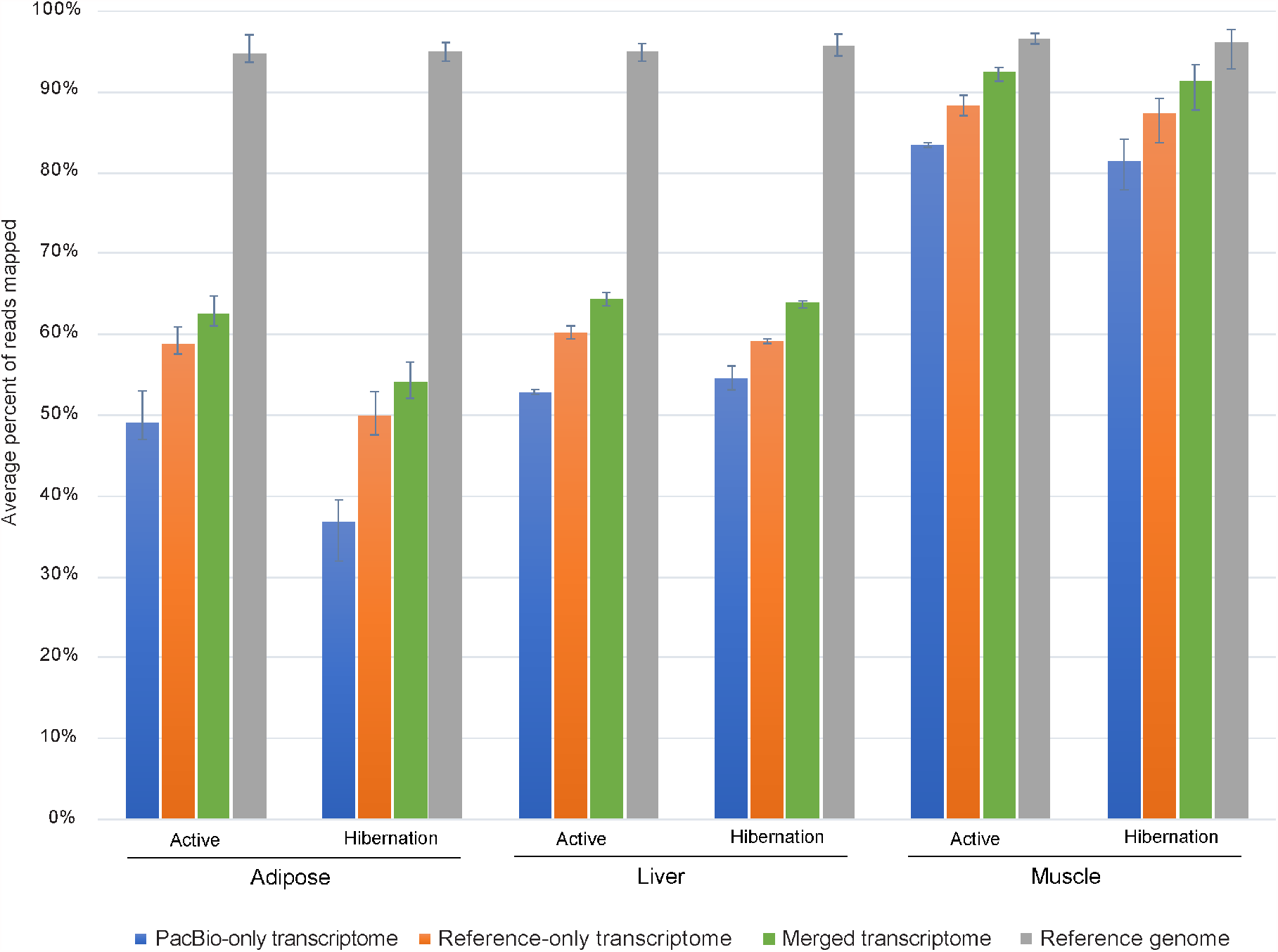
Ribosomal RNA depleted RNA-seq data mapped to transcriptomes and the reference genome. Average percent of reads mapped per season and tissue. Error bars indicate range.

### Tissue-specific Differential Isoform Usage from Active to Hibernation Season

We assessed whether the relative abundance of different isoforms for each gene varied between the seasons (differential isoform usage, DIU). We also determined whether the major (highest expressed) isoform switched between seasons. When analyzing for DIU and major isoform switching between the seasons (hibernation vs active), there were substantial differences among the tissues (Table 1, Figure S3). Adipose had the highest incidence of DIU with regards to both number and percent of all analyzed genes, with and without major isoform switching (27.5% of genes; Table 1). In contrast, major isoform switching between states without differential isoform usage was fairly consistent across tissues.

**Table 1:** Number and percentage of genes showing differential isoform usage (DIU) and/or major isoform switching between hibernation and active seasons in each tissue.

|  | **Adipose** | **Liver** | **Muscle** |
| --- | --- | --- | --- |
| DIU – Major Isoform Switching | 779 (8.7%) | 201 (2.7%) | 85 (1.6%) |
| DIU – No Major Isoform Switching | 1679 (18.8%) | 379 (5.0%) | 198 (3.7%) |
| Not DIU – Major Isoform Switching | 1196 (13.4%) | 1228 (16.3%) | 826 (15.4%) |
| Not DIU – No Major Isoform Switching | 5268 (59.0%) | 5706 (75.9%) | 4255 (79.3%) |
| Total Genes Analyzed | 8922 | 7514 | 5364 |

### Isoform Switching between Active and Hibernation Seasons Despite No Gene-Level Expression Change

In this study and in prior work, gene expression changes were detected in all three tissues between active and hibernating seasons. With the new isoform-level quantifications, we examined whether the genes classified as having both DIU and Major Isoform Switching between the two seasons (Table 1) were enriched in specific cellular functions. Of the three tissues, adipose displayed the most DIU + Major Isoform switches between seasons, with the encoded genes displaying enrichment for biosynthetic and metabolic processes (Table S4). We then restricted the list of DIU and Major Isoform switching genes to those that displayed less than 20% change in overall gene-level expression in response to the season. Testing this list of genes for enrichments in cellular function found a shift to more modest enrichments with the top categories focused on autophagy (Table S5). This indicates that one component of hibernation adaptations, the need to consume reserve fat stores, is partly re-programmed by a shift in mRNA isoform ratios rather than the overall expression level of the gene.

One gene that shows a dramatic isoform switch in adipose in concert with the seasons is Integrin Subunit Beta 3 Binding Protein (*ITGB3BP*), imparting a nearly binary switch in five of the six sampled bears (Figure 4). There is very little change in overall gene expression but the major isoform switches between active and hibernation (Figure 4a, b). The two isoforms differ by a cassette exon near the 3’ end of the open reading frame, resulting in a new C-terminal peptide sequence for the protein. Prior characterization of the ITGB3BP protein in mammals demonstrated that it acts as transcriptional co-regulator amongst several nuclear hormone receptor circuits affecting both the retinoic acid (RXR) and thyroid hormone (TR) pathways. Since nuclear hormones play a key role in metabolic control and are implicated in homeostasis during hibernation, we further investigated the putative consequences of the isoform switch [23]. The bear isoform PB.6860.1, which is predominant in bear adipose tissue during hibernation, includes a cassette exon near the 3’ end of the open reading frame and encodes a protein isoform of the same 177 amino acid length as the predominant human isoform with 76% identity (Figure 4d). Skipping of the cassette exon (PB.6860.2) is more common during the active season, and this splicing pattern is also observed in humans, according to GTeX project data [24]. This event results in a truncated C-terminus, which we predict would alter the downstream co-regulator activity of ITGB3BP based on studies of its domain structure. Studies of the human protein showed that the C-terminal LXXIL motif, conserved in the bear and encoded by the exon-included isoform (PB.6860.1), acts as a receptor-interaction domain (RID) [25]. This motif along with the human N-terminal LXXLL motif, also conserved in the bear, act in a cooperative fashion during interaction with the nuclear hormone receptor dimer. As the isoform without the exon (PB.6860.2) lacks the C-terminal LXXIL motif and terminates instead with a dipeptide GI, this alternative splicing event may impart broad downstream consequences for hormone receptor signaling in adipose tissue while bears are in an active state.

**Figure 4.**
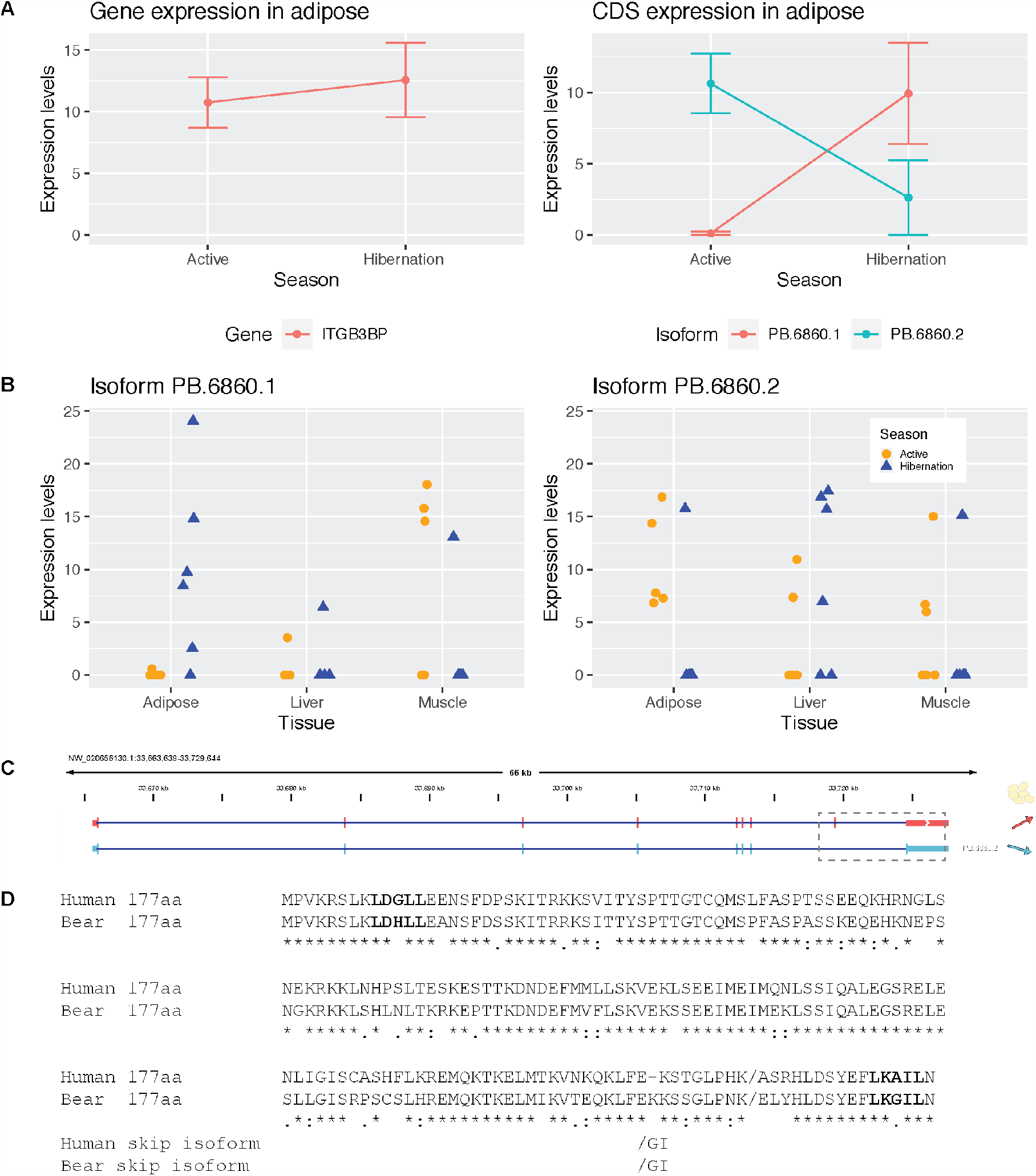
Differential isoform expression of *ITGB3BP* mRNA in adipose tissue. (a) expression changes at the gene and coding sequence (CDS) level. (b) Short read expression of the two *ITGB3BP* isoforms across all bears and tissues. (c) Isoform structure of two *ITGB3BP* isoforms, dashed box indicates the region of the isoforms that differs. (d) Protein alignment (MUSCLE) for the human isoform and the bear isoform observed in the active season. Putative functional motifs based on the human protein co-activator function are shown in larger bold text. The bold slash represents the beginning of C-terminal peptide sequence altered by the splicing event observed in hibernating bears. The skipped exon results in a shorter C-terminus shown below the full-length alignments.

In liver, an example of a gene with significant DIU and isoform switching was Carbonic Anhydrase 2 (*CA2*). The *CA2* gene expressed two isoforms (Figure S4). The shorter isoform, which lacks a coding exon and initiates transcription downstream compared to the other isoform, was upregulated in the hibernation season for all six bears, while the longer isoform showed decreased expression during hibernation. Interestingly, the shorter isoform, which is annotated in the human genome, was not annotated in the reference transcriptome and was only found in the PacBio Iso-Seq data (Figure S1e).

In muscle, an example of a gene with significant DIU and isoform switching, was cryptochrome 2 (*CRY2*). There are four expressed isoforms of *CRY2*, two of which are novel and showed very little changes from active season to hibernation and the other two isoforms, which show major isoform switching (Figure S5). The two isoforms that show major isoform switching differed by only 14 bp at the last donor site; both isoforms were annotated in the reference, and both were predicted to encode for the same protein. The longer isoform showed decreased isoform usage while the shorter isoform showed increased isoform usage during hibernation.

In addition to identifying genes with DIU and major isoform switching, we also identified genes with evidence for DIU without major isoform switching (Table 1). For example, four transcript isoforms of the homeobox transcription factor *PRRX1* (Figure S6), corresponding to human *PRRX1a* (PB.14470.3); *PRRX1b* (PB.14470.1), and an unnamed human isoform (PB.14470.4). The fourth transcript, PB.14470.2, is a non-coding isoform. Transcription factor *PRRX1* is highly expressed in adipose and muscle. In adipose, the PB.14470.1 isoform continues to predominate in both active and hibernation seasons compared to the other isoforms, but also shows significantly higher expression during hibernation. This isoform (PB.14470.1) also contains a 3280 bp 5’ UTR that is lacking in PB.14470.3 while the latter contains a fifth exon. This same transcription factor in muscle shows no change in isoform expression between the seasons (Figure S6).

A gene that shows DIU and is positively regulated in adipose by *PRRX1* in humans is COL6A3. *COL6A3* expression is positively related to *PRRX1* in hibernation, but not in active season adipose (Figure S7); overall the relationship between these two genes is opposite to that in humans. Two coding isoforms (out of six) of *COL6A3* predominate and both isoforms are reduced in hibernation compared to active season (Figure S7).

## Discussion

Our study provides an unprecedented view into hibernation biology through the lens of RNA processing by producing a dataset that improved the annotation of the brown bear genome and reinforced the important role adipose tissue plays in hibernation. This approach allowed us to characterize isoforms that are changing between active and hibernation states, even when the gene itself shows no significant change in expression levels. While studies of differentially expressed genes have provided much of our current understanding in hibernation biology [15, 26-31], determining genes where functionally distinct isoforms change between seasons is the next essential biological mechanism to uncover.

As we have shown in this study, metabolically active tissues vary dramatically in their isoform usage. The inherent complexity and importance of adipose, which has been recognized as a source of critical physiological mechanisms during hibernation [7, 15], continues to expand. The large percentage of genes with DIU compared to liver and muscle, and the large number of differentially expressed genes support the dynamic role of adipose in hibernation where both transcription and RNA processing play concerted roles. The difference in the percentage of reads mapping to the transcriptome as compared to the genome in adipose and liver suggests that the ribosomal-depleted RNA-seq data had more intronic and intergenic reads in those tissues. It is also important to note that slightly different extraction kits were used for the muscle as compared to adipose and liver tissue RNA extraction for the short-read data [15]. While extraction method may influence the percentage of intronic and intergenic reads, there may also be differences in splicing efficiency between tissues and states. Regardless, adipose showed differences between active and hibernation, suggesting that intron retention may be an especially important mechanism for gene regulation and function in adipose during hibernation. It is also possible that global rates and efficiency of RNA processing by the spliceosome may also vary between the tissues and states.

In addition to hibernation biology, our results also highlight several transcriptional events that may have important implications for humans. For example, the very strong positive relationship between *COL6A3* and *PRXX1* expression in human adipose tissue [32] is completely absent in bears, providing a possible explanation for why bears exhibit little to no inflammatory signatures even during periods of extreme adiposity and when consuming a high saturated fat diet [33]. Even the expected (positive) relationship between *COL6A3* and TGFß is absent in bears. However, elevated PRRX1 is associated with PPARG (Peroxisome Proliferator Activated Receptor Gamma) reductions in hibernation bear adipose tissue as has been observed previously [34]. Thus, certain aspects of adipose function appear dissociable in bears which could lead to a more precise understanding of the contribution of adipose tissue to human disease states. Furthermore, since various single nucleotide polymorphisms within the homeobox domain superfamily associate with type 2 diabetes in humans, manipulation of PRRX1 isoforms in hibernation bear adipocytes (insulin resistant) *in vitro* could be used to test these relationships more precisely without the confounding interaction with COL6A3.

While PacBio Iso-Seq has been used widely for plant and animal genome annotation [35, 36], and recently shown to shed light in cell-type specific isoform expressions in brain regions [37], this study included multiple biological replicates per tissue and condition that incorporated both RNA-seq and Iso-Seq data for identifying differential isoform usage. The high mappability of the matching RNA-seq data to the Iso-Seq transcriptome shows promise for other less well-annotated organisms, where Iso-Seq may serve as a sole reference transcriptome upon which RNA-seq may be used for differential analysis. We saw a high correlation at the gene level between data types from the same tissue despite the fact that the Iso-Seq data were lower coverage. There was also a moderate correlation at the isoform level within data type, but less so between data types, likely because of the low sequencing depth of the long-read data.

Moving forward, we aim to create an improved reference transcriptome at higher coverage, as well as improve the existing reference genome, to create a comprehensive reference that may better serve for studying differential isoform expressions in hibernation. Comparative studies across bears will also provide a much-needed framework for comparing hibernating and non-hibernating species. Indeed, long-read sequencing was used to improve the polar bear reference genome annotation set [38], which provides a valuable additional resource for comparative studies. Although the *U. arctos* genome [39] is not a chromosome-level assembly, we were able to improve the annotations and these data can be used in future improvements of the genome assembly.

In summary, our study demonstrates the utility of PacBio Iso-Seq for determining isoform differences between hibernating and active brown bears. Importantly, we found that adipose is the most dynamic tissue during hibernation with the highest number of genes with differential isoform usage and isoform switching as compared to liver and muscle. These findings and datasets provide a rich new resource for studying hibernation biology and understanding metabolic function. Additionally, this resource provides a basis for incorporating isoform changes in studying hibernation and translating findings to solving human diseases.

## Methods

### Sample Collection

For the PacBio Iso-Seq protocol, samples were collected from three bears (1 female, 2 males) at the Washington State University Bear Research, Education, and Conservation Center in January 2019 and May 2019, to represent the winter hibernation and summer active periods, respectively. Muscle, liver, and adipose tissue samples were collected for a total of 18 samples. Samples for the Illumina Ribo-Zero RNA-seq data were collected in May 2015 and January 2016 and are described in detail in [15]. Animal care details are described in [7]. Bears were first anesthetized using the protocol described in Ware JV, Nelson OL, Robbins CT and Jansen HT [40]. Subcutaneous adipose samples were collected using a 6mm punch biopsy (Miltex, York, PA) as described in [7], while skeletal muscle (gastrocnemius) and liver tissue samples were collected with a 14G tru-cut biopsy needle (Progressive Medical International, Vista, CA, USA). All samples were collected between 0800 and 1200hr. Samples were immediately flash frozen in liquid nitrogen and transferred to a -80°C freezer, where they were stored until shipment to the University of Delaware Sequencing & Genotyping Center for RNA extraction. Procedures for all experiments were approved by the Institutional Animal Care and Use Committee at Washington State University (Protocol #04922).

### PacBio Iso-Seq Library Preparation and Sequencing

Total RNAs were extracted from tissue samples and isolated using the RNeasy Universal kit (Qiagen, Valencia, CA, USA) as per the manufacturer’s standard protocol. Following total RNA isolation, the samples were concentrated using RNA Clean & Concentrator Kit (Zymo Research, Irvine CA, USA). The purity of RNA samples was measured using the DeNovix DS-11+ spectrophotometer (DeNovix Inc., Wilmington, DE, USA). RNA concentration was measured using Qubit High Sensitivity RNA Assay Kit and Qubit 3.0 Fluorometer (Thermo Fisher Scientific Inc., Waltham, MA, USA). The integrity of total RNA was assessed on the Agilent Fragment Analyzer 5200 system (Agilent Technologies, Santa Clara, CA, USA) using the High Sensitivity RNA Kit. The RNA Quality Number (RQN) criteria for the RNA samples was RQN >7.0.

From 100ng to 300ng of total RNA was input for cDNA synthesis and amplification using NEBNext Single Cell/Low Input cDNA Synthesis & Amplification Module (New England BioLabs Inc., Ipswich, MA, USA) as per the manufacturer’s standard protocol. This is a poly-A selection library preparation method. A total of 10-15 PCR cycles were used to generate sufficient quantities of cDNA for PacBio Iso-Seq library preparations. Concentration and size profile of cDNA samples was assessed on the Agilent Fragment Analyzer 5200 system (Agilent Technologies, Santa Clara, CA, USA) using the High Sensitivity Large Fragment Kit.

Amplified cDNA samples were size selected using ProNex Size-Selective Purification System (Promega Corporation, Madison, WI, USA) as per the PacBio recommendation for standard length cDNA transcripts. Size selected cDNA was used to construct SMRTbell Iso-Seq libraries using Express Template Prep 2.0 (Pacific Biosciences, Menlo Park, CA, USA) as per the manufacturer’s Iso-Seq Express Template Preparation protocol. The concentration of the Iso-Seq libraries was measured using the Qubit 3.0 Fluorometer (Thermo Fisher Scientific Inc., Waltham, MA, USA). The fragment size profile of the Iso-Seq libraries was assessed on the (Agilent Technologies, Santa Clara, CA, USA). Each Iso-Seq library was run on a single Sequel system SMRT Cell using sequencing chemistry 3.0 with 4 hour pre-extension and 20 hour movie time. One SMRT Cell per tissue and state was used to provide deep coverage of the entire grizzly transcriptome. Raw reads were processed into circular consensus sequence (CCS) reads as per the manufacturer’s standard pipeline (SMRT Link version 7.0).

### PacBio Iso-Seq Bioinformatic Analysis

All 18 SMRT Cells of Iso-Seq data from different samples were pooled and run through the IsoSeq Analysis in SMRTLink v8.1 which generated full-length, high-quality (HQ) isoform sequences. The HQ isoforms were mapped to the genome assembly (GCA_003584765.1) using minimap2, then filtered and collapsed into non-redundant isoforms using Cupcake following the analysis described at (https://github.com/Magdoll/cDNA_Cupcake/wiki/Cupcake:-supporting-scripts-for-Iso-Seq-after-clustering-step). The non-redundant isoforms were then classified against the GCA_003584765.1 reference transcriptome using SQANTI3 (https://github.com/ConesaLab/SQANTI3). After running SQANTI3 classification and filtering, we obtained a final set of PacBio isoforms that we used subsequently as the reference transcriptome for short read quantification. As part of the Cupcake processing pipeline, we obtained full-length read counts associated with each isoform, which were then normalized into Full-Length Counts Per Million (CPM) for cross-sample comparison.

### Short-read RNA-seq Quantification

Short-read Illumina data from [15] for the same individuals and tissues, sampled in a different year, were mapped to the final set of PacBio isoforms using the kallisto quantification algorithm, with the rf-stranded flag [43]. Read counts were compared to the IsoSeq data using Pearson correlation on the log_2_ count per million.

### Merging annotations and mapping

The PacBio-only annotation set was merged with the reference-only annotations using gffcompare [41]. Short read Illumina data from [15] was mapped to each transcriptome and the reference genome (GenBank assembly accession: GCA_003584765.1 [39]) using HISAT2 (version 2.2.1) with the --rf flag and otherwise default parameters [42].

### Functional Annotation and Differential Isoform Expression Analysis

Abundance count estimates for each individual were combined into a single input matrix for tappAS [44]. The annotation file generated in SQANTI3 and the short-read count matrices for each tissue were input into tappAS [44]. Each tissue was analyzed separately to compare differential isoform expression in hibernation compared to active season. We excluded short-read data from sample 1AA because it was previously shown to contain a hair follicle [15]. Transcripts with counts per million (cpm) less than 1.0 or coefficient of variation cutoff of 100% were excluded from the analyses. In the differential isoform usage (DIU) analysis, minor isoforms with a proportional of expression difference less than 0.1 were excluded from the analysis and DIU was considered at an FDR < 0.05. Gene ontology (GO) enrichment of different gene sets was calculated using PANTHER [45] with the following parameters: Analysis

Type: PANTHER Overrepresentation Test (Released 20210224), Annotation Version and Release Date: GO Ontology database DOI: 10.5281/zenodo.4495804 Released 2021-02-01, Reference List: Homo sapiens (all genes in database), Annotation Data Set: GO biological process, Test Type: Fisher’s Exact, Correction: Calculate False Discovery Rate.

## Declarations

### Ethics Approval and Consent to Participate

The bears used in this study were housed at the Washington State University Bear Research, Education, and Conservation Center. All procedures were approved by the Washington State University Institutional Animal Care and Use Committee (IACUC) under protocol number ASAF 6546.

### Availability of Data and Materials

The datasets generated for the current study are available in the NCBI SRA repository under BioProject PRJNA727613. The datasets reanalyzed in this study are available in the NCBI SRA repository under BioProject PRJNA413091. The code for this project is available at: https://github.com/jokelley/brownbear-isoseq-act-hib

## Authors’ Contributions

BDEH, ST, BK, MV, CTR, HTJ, JLK contributed to sampling. BK, OS, and EB performed the RNA extraction, library preparation, and sequencing. JGU, MV, and HTJ contributed to the manuscript. ET and JLK analyzed the data and wrote the manuscript. All authors read and approved the final manuscript.

## Acknowledgements

This research used resources from the Center for Institutional Research Computing at Washington State University. Allan Cornejo Kelley for feedback on figures.

